# Functional selectivity and specific connectivity of inhibitory neurons in primary visual cortex

**DOI:** 10.1101/294835

**Authors:** Petr Znamenskiy, Mean-Hwan Kim, Dylan R. Muir, Maria Florencia Iacaruso, Sonja B. Hofer, Thomas D. Mrsic-Flogel

**Affiliations:** Biozentrum, University of Basel, Basel, Switzerland; Sainsbury Wellcome Centre, UCL, London, UK; Allen Institute for Brain Science, Seattle, USA; aiCTX AG, Zurich, Switzerland; University of Oxford, Oxford, UK

## Abstract

In the cerebral cortex, the interaction of excitatory and inhibitory synaptic inputs shapes the responses of neurons to sensory stimuli, stabilizes network dynamics^1^ and improves the efficiency and robustness of the neural code^2–4^. Excitatory neurons receive inhibitory inputs that track excitation^5–8^. However, how this co-tuning of excitation and inhibition is achieved by cortical circuits is unclear, since inhibitory interneurons are thought to pool the inputs of nearby excitatory cells and provide them with non-specific inhibition proportional to the activity of the local network^9–13^. Here we show that although parvalbumin-expressing (PV) inhibitory cells in mouse primary visual cortex make connections with the majority of nearby pyramidal cells, the strength of their synaptic connections is structured according to the similarity of the cells’ responses. Individual PV cells strongly inhibit those pyramidal cells that provide them with strong excitation and share their visual selectivity. This fine-tuning of synaptic weights supports co-tuning of inhibitory and excitatory inputs onto individual pyramidal cells despite dense connectivity between inhibitory and excitatory neurons. Our results indicate that individual PV cells are preferentially integrated into subnetworks of inter-connected, co-tuned pyramidal cells, stabilising their recurrent dynamics. Conversely, weak but dense inhibitory connectivity between subnetworks is sufficient to support competition between them, de-correlating their output. We suggest that the history and structure of correlated firing adjusts the weights of both inhibitory and excitatory connections, supporting stable amplification and selective recruitment of cortical subnetworks.

---

Cortical neurons derive their functional properties from the patterns of their synaptic inputs. Thus a characterization of the rules that govern these patterns is essential for understanding the mechanisms of cortical computations. While pyramidal neurons in the cortex are organized into specific excitatory subnetworks and make connection with only a small fraction of nearby excitatory cells that receive common feedforward inputs^14^ and share their response properties^15,16^, inhibitory neurons, such as parvalbumin-expressing (PV) basket cells, are thought to make dense and unstructured connections within cortical networks^9–12^. Although some studies have challenged this conclusion^17–19^, the prevailing view is that PV neurons pool their inputs and provide uniform inhibition to the surrounding excitatory neurons.

The dense nonspecific connectivity of PV neurons is seemingly at odds with the observation that inhibitory inputs onto individual pyramidal cells, while often broadly tuned, tend to match the tuning preference and strength of excitatory inputs^6,20^ even for features that are not organized in a columnar fashion^7,8,21^. However, these observations can be reconciled if the inhibitory interneurons reflect the diversity of the local excitatory population rather than merely its average and provide stronger inputs to contemporaneously active pyramidal cells^22^. To test these predictions, we simultaneously characterized visual responses and synaptic connectivity of PV-positive interneurons and pyramidal cells in mouse primary visual cortex.

We expressed GCaMP6f in transgenic mice expressing tdTomato in PV-positive interneurons and recorded the visual responses of PV-positive and PV-negative cells in layer 2/3 of primary visual cortex using two-photon calcium imaging. We used a rich stimulus set including full field gratings of varying direction, spatial and temporal frequency (Figure 1a). We quantified the tuning selectivity of single neurons by measuring the skewness of the distribution of responses over the set of visual stimuli. PV neurons tended to be less selective than pyramidal cells, responding to a broader range of stimuli (Figure 1b; *p* = 8.8×10^−56^, ranksum test, 209 PV+ and 7399 PV- neurons from 10 mice). In agreement with previous reports^10,12^, PV cells were broadly tuned to orientation and direction, as well as spatial and temporal frequency (Figure 1c–f). Furthermore, spatial and temporal frequency tuning of PV cells was often multi-peaked and poorly structured (e.g. Figure 1k, cell 1).

**Figure 1:**
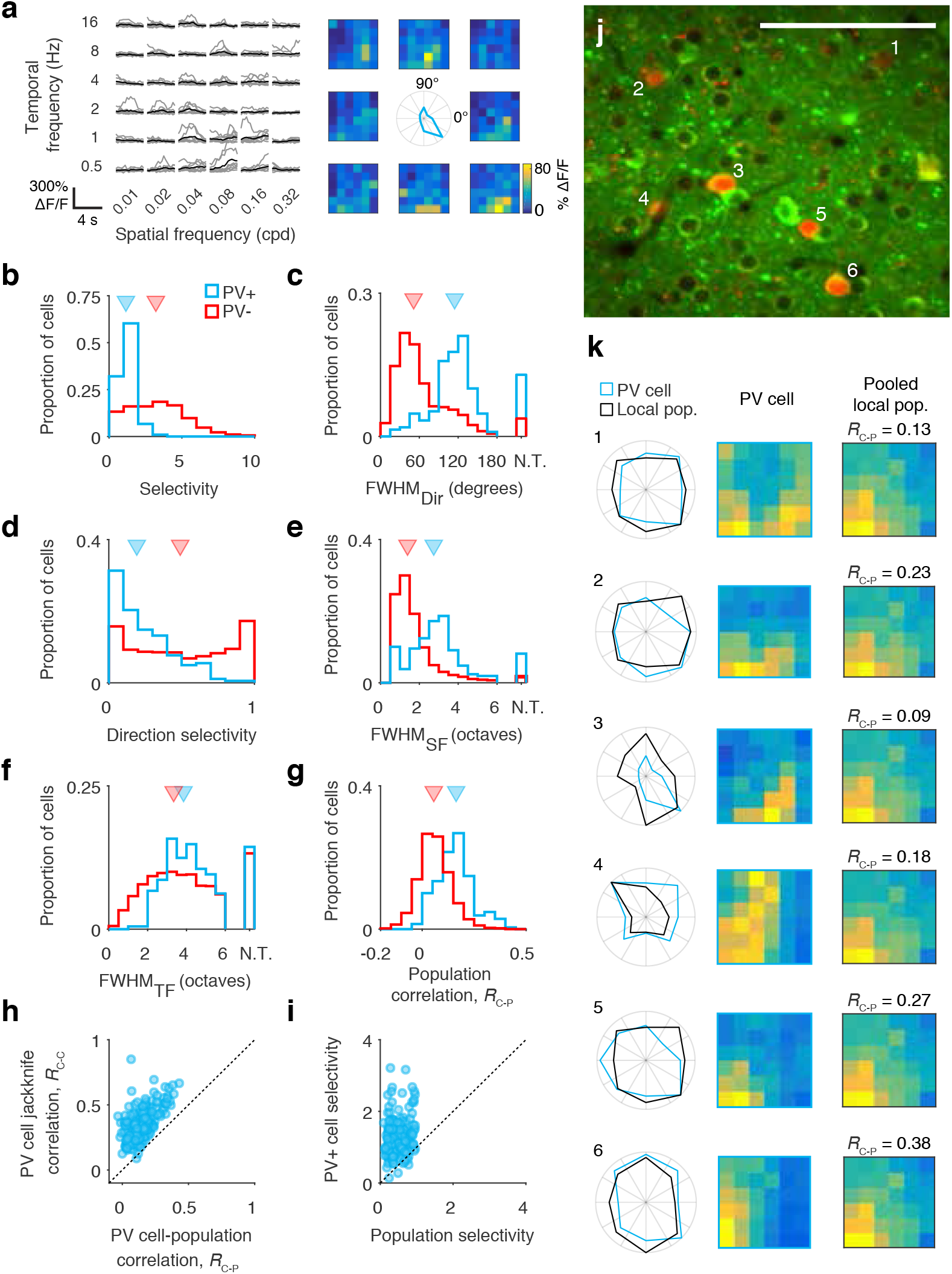
Response properties of PV neurons in layer 2/3 of mouse visual cortex. **a**. Responses of an example PV neuron to sinusoidal gratings of different spatial and temporal frequencies. Traces show individual (gray) and mean (black) responses of the cell at its preferred direction (315 degrees). Color maps show mean responses to the moving phase of the grating across all directions. **b**. PV+ cells are less selective than PV- cells, quantified by skewness of their F/F distributions. Triangles indicate medians. **c**. PV+ cells are broadly tuned to orientation. N.T.: cell untuned for orientation. **d**. PV+ cells are less direction selective than PV- cells. **e–f**. PV+ cells are more broadly tuned to spatial (**e**) and temporal (**f**) frequency than PV- cells. **h**. Local population activity predicts the responses of PV+ cells better than that of PV- cells. **i**. PV+ cells tend to be more selective than local population activity. **j**. Locations of PV+ cells depicted in panel k within the field of view. Note that some of the cells were recorded in a different imaging plane. Scale bar: 100 *µ*m. **k**. Comparison of PV cell responses to the local population. Polar plots show responses at spatial and temporal frequency that evoked the highest mean response (blue: PV cell; black: local population). Colormaps show spatial and temporal frequency responses averaged across direction. *R*_*C-P*_ values quantify the correlation between single cell responses and population mean as in panel **g**.

According to the non-specific pooling model, the tuning of individual PV neurons reflects the average selectivity of nearby pyramidal cells^12,13^. To determine the extent to which visual responses of PV neurons could be explained by local biases in tuning, we attempted to predict the tuning of individual cells by computing a weighted average of the responses of surrounding neurons, with Gaussian drop-off of weights with distance. Single-trial responses of individual cells were predicted by computing the local population activity averaged over all other trials. In agreement with previous reports^12^, pooled local population activity was typically better at predicting the selectivity of PV neurons than that of pyramidal cells (Figure 1g, *p* = 9.9 × 10^−53^, ranksum test). However, the predictions were typically poor even for PV cells (median correlation between PV cell responses and population prediction *R*_*C-P*_ = 0.16).

**Supplemental Figure 1:**
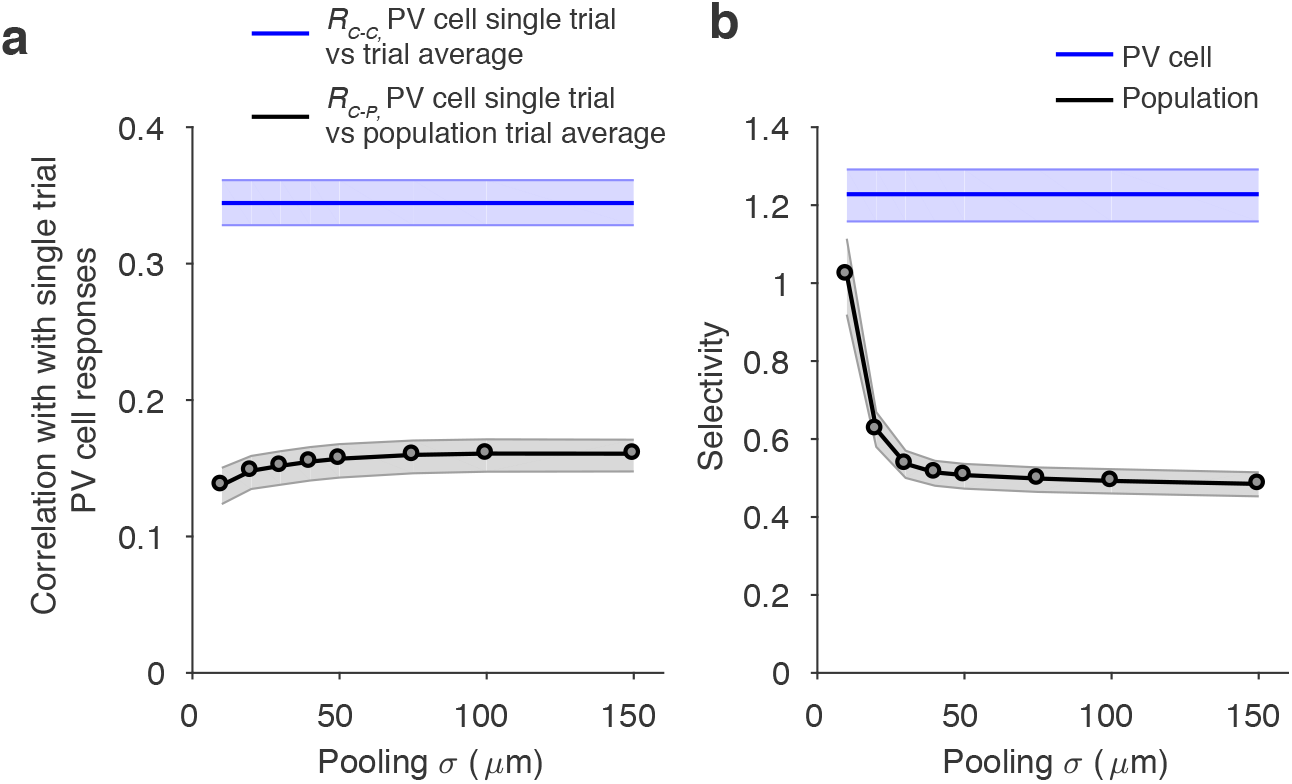
Pooled population activity falls short of predicting PV neuron visual selectivity, independent of the spatial scale of pooling. **a**. Mean correlation of single trial PV cell responses with local population (black) or same cells response (blue) averaged across all other trials for different spatial scales of population pooling. Shading 95% confidence interval, derived by bootstrap resampling across cells. **b**. Selectivity of PV cell and local population responses across for different spatial scales of population pooling. Shading 95% confidence interval, derived by bootstrap resampling across cells.

Do the weak correlations between single PV cell and population responses reflect high response variability, or a true deviation from the non-specific pooling model? We estimated the upper bound for response predictions imposed by measurement noise and trial-to-trial neural variability by computing the correlation between single trial visual responses and the cell’s own mean response on other trials. This estimate consistently outperformed the population prediction (Figure 1h), indicating that individual PV cell responses indeed deviate significantly from the local population. These deviations were apparent in the direction, spatial and temporal frequency tuning profiles of individual PV cells, which often differed dramatically from the local population and from other nearby interneurons (Figure 1j–k). Finally, responses of PV cells were consistently more selective than the population mean response (Figure 1i). These conclusions were independent of the spatial scale of population pooling (Supplemental Fig. 1). Thus, responses of PV neurons are heterogeneous and not fully explained by biases in the tuning of the local population.

To understand the origin and impact of the diverse responses of PV neurons within cortical microcircuits, we examined the organization of PV cell connections with pyramidal cells using whole-cell patch clamp recordings in acute slices. Consistent with previous reports, we found that PV cells were reciprocally connected with the majority of nearby excitatory neurons (Figure 2a; 157 PV to Pyr pairs, 138 Pyr to PV pairs from 35 mice). However, the strengths of both excitatory inputs onto PV cells and their inhibitory inputs onto pyramidal cells spanned almost two orders of magnitude and followed a log-normal distribution (Figure 2b–c). In reciprocally connected pairs the magnitude of the EPSP onto the PV neuron was highly correlated with the magnitude of the IPSP onto the pyramidal cell (Figure 2d–e, 88 connections including 58 PV cells and 79 pyramidal cells from 29 mice). This relationship persisted when controlling for distance between patched cells (partial correlation *R* = 0.57, *p* = 1.2×10^−8^; Supplemental Fig. 2). To exclude the possibility that this correlation arose as a consequence of variability in slice quality, we examined recordings where we simultaneously measured the strength of connections between a PV cell and multiple pyramidal cells. We normalized the strength of excitatory and inhibitory connections by the geometric mean EPSP or IPSP magnitude of all simultaneously recorded cells. The correlation between the strength of inhibitory and excitatory connectivity persisted following this correction for variation in slice quality (Figure 2f; 53 pairs including 23 PV cells from 16 mice; *p* = 0.0095, see methods). Together, these observations show that the strength of excitatory and inhibitory connections of PV interneurons is heterogeneous and organized in a non-random manner. Contrary to the nonspecific pooling model, individual PV cells receive the strongest excitatory inputs from a subset of nearby excitatory neurons, to which they provide strong feedback inhibition.

**Figure 2:**
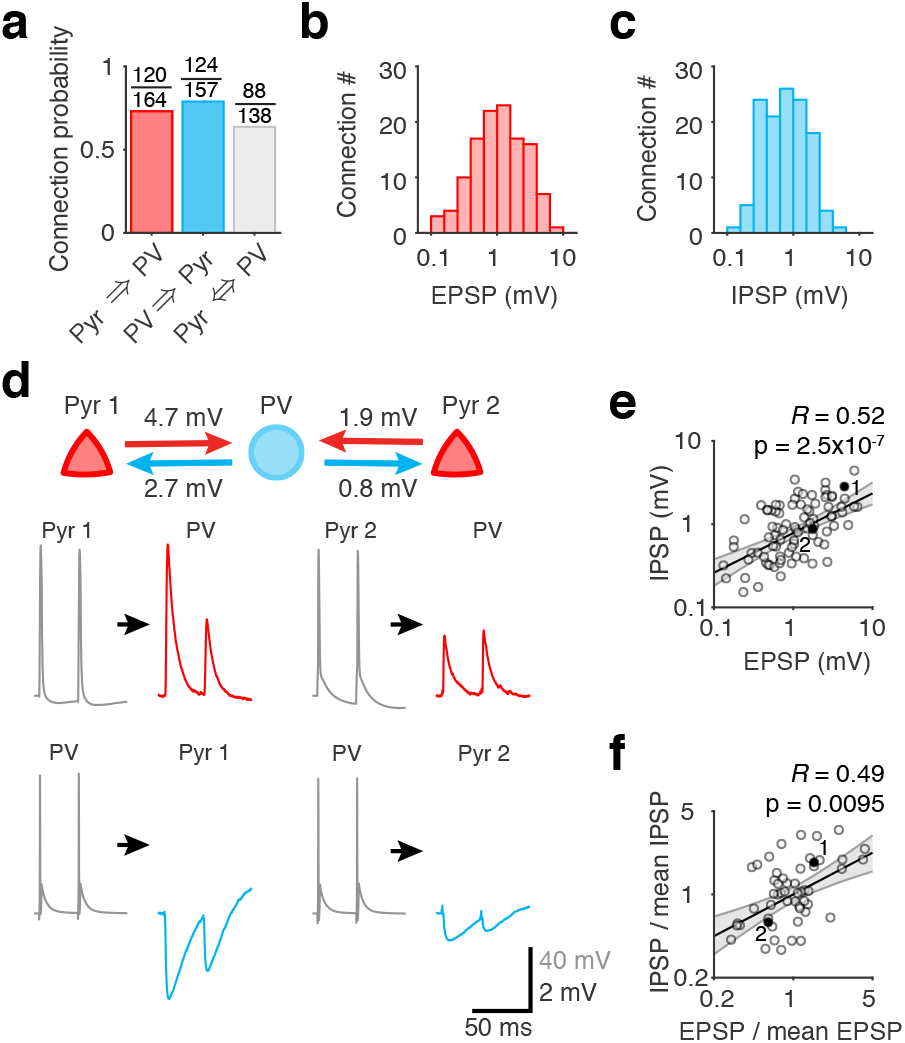
Heterogeneity of synaptic strength of PV neuron connections. **a**. Rates of pyramidal / PV neuron connectivity. **b.** Distribution of the strength of excitatory connections from pyramidal cells onto PV neurons. **c.** Distribution of the strength of inhibitory connections from PV neurons onto pyramidal cells. **d.** Example recording of a PV neuron reciprocally connected to two pyramidal cells. PV neuron provides stronger inhibition to the pyramidal cell that provides it with stronger excitatory input. **e.** EPSP and IPSP strengths are correlated for reciprocally connection PV / pyramidal neuron pairs. Black line: best fit regression line of log-IPSP magnitude against log-EPSP; gray shading: 95% confidence interval for the regression line estimated from bootstrap resampling; R and p are Pearson correlation and its p-value, respectively. Cell pairs in panel **d** are highlighted. **f.** In recordings with multiple pyramidal neurons reciprocally connected to a PV cell, the correlation between EPSP and IPSP magnitude persists after controlling for slice quality by normalizing each by the geometric mean of EPSP/IPSP strength in the recording. Notation as in panel **e.** P-value was estimated using a shuffling procedure (see methods).

**Supplemental Figure 2:**
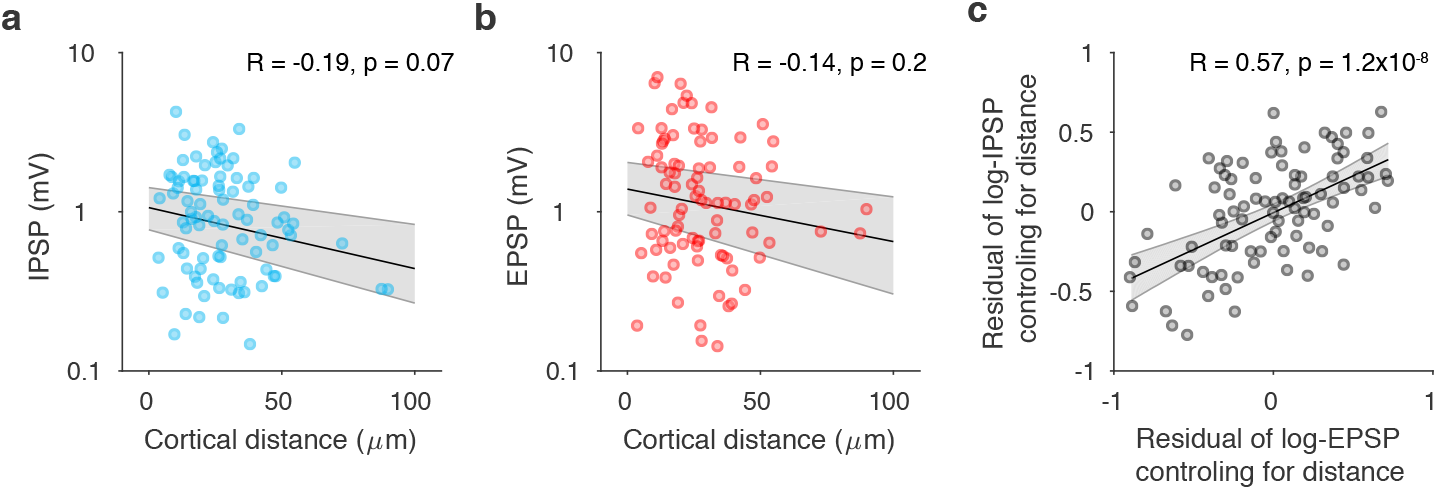
Distance does not account for the correlation between EPSP and IPSP magnitude for reciprocally connected PV / pyramidal cell pairs.

To determine how the heterogeneity of PV neuron synaptic weights relates to their visual responses, we identified pyramidal and PV cell pairs, whose visual responses we characterized *in vivo*^10,15,16^ (Figure 3a–c). We first examined the relationship between responses of PV/pyramidal neuron pairs and their connection probabilities, using a cosine similarity measure to quantify overlap in their activity patterns (total response similarity). Consistent with previous reports, we found that PV neurons provided inhibition to and received excitatory input from the majority of nearby excitatory cells, independent of the similarity of their responses (Figure 3d–e; 56 PV to Pyr pairs from 21 mice, 55 Pyr to PV pairs from 20 mice). Since absolute rates of connectivity are difficult to estimate in vitro due to artefacts introduced by the slicing procedure, the absence of a relationship between PV / pyramidal connection probability and functional similarity is consistent with the interpretation that PV cells make connections with all or nearly all nearby pyramidal cells, independent of their functional properties^9^.

We next examined the strength of synaptic connections of PV neurons. In contrast to connection probability, the magnitude of IPSPs from PV cells onto pyramidal cells was positively correlated with the similarity of their activity patterns (Figure 3f; 46 pairs from 18 mice, *R* = 0.55, *p* = 8×10^−5^). The strength of excitatory connections from pyramidal cells onto PV cells showed a similar but weaker trend (Figure 3g; 37 pairs from 19 mice, *R* = 0.33, *p* = 0.044). To determine whether this relationship was driven by the similarity of the cells’ visual tuning, we computed response similarity between trial-averaged response traces from pairs of cells, and related this metric to the strength of synaptic connections between the pair. This signal response similarity measure was significantly correlated with the strength of connections formed by PV cells with excitatory neurons (Figure 3h–i, Supplemental Fig. 3–5). This correlation was not observed if we discarded information about the time-course of the cells responses or examined the similarity of direction, spatial, or temporal frequency tuning in isolation (Figure 3h–i, Supplemental Figure 4–6). On the contrary, just as for connections between pairs of excitatory neurons^15^, connections between PV neurons and pyramidal cells appear to adjust their synaptic weights based on the overall similarity of the responses of the pre- and post-synaptic cell, consistent with Hebbian plasticity of inhibitory connections^22^.

**Figure 3:**
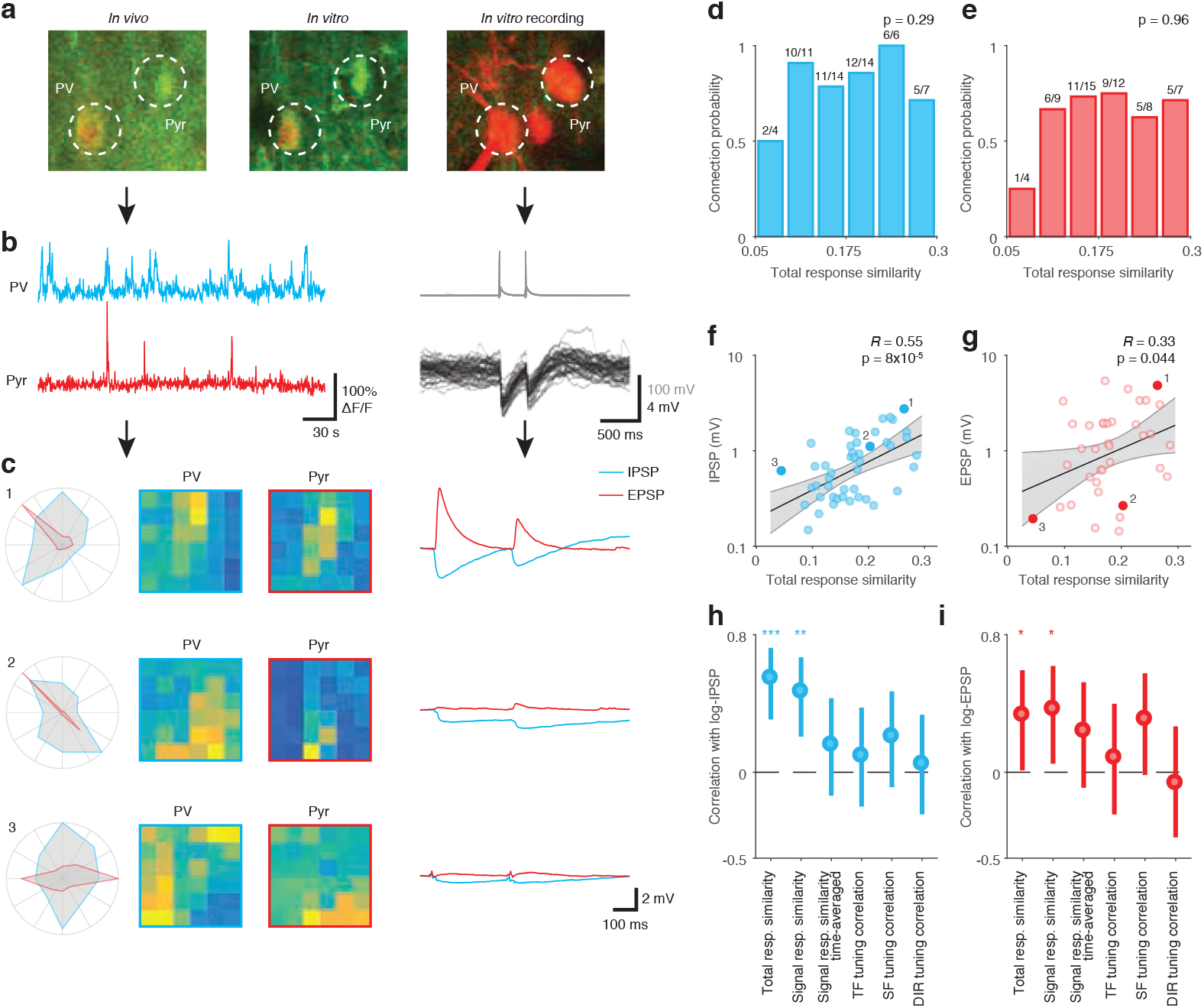
Functional similarity predicts the strength of PV neuron connections. **a.** PV-pyramidal neuron pair identified during in vivo imaging (top left), and *in vitro* recording (top middle and right). The in vivo image was generated by resampling the in vivo Z-stack to match to the coordinate frame of the *in vitro* volume. **b.** Example in vivo calcium traces (left) of the cells in panel a and *in vitro* current-clamp recordings of evoked action potentials in the PV cell (top right) and IPSPs in the pyramidal cell (bottom left). **c.** Visual tuning and postsynaptic potentials for 3 reciprocally connected PV / pyramidal cell pairs. Cell pair 1 is depicted in panels **a–b. d.** Frequency of inhibitory connections from PV onto pyramidal cells does not depend on total response similarity. P-value corresponds to the slope coefficient of logistic regression of connection probability against total response similarity. **e.** Frequency of excitatory connections from pyramidal onto PV cells does not depend on total response similarity. **f.** Strength of inhibitory connections from PV onto pyramidal cells correlates with total response similarity. Cell pairs shown in panel c are highlighted. Black line: best fit regression line of log-IPSP magnitude against response similarity; gray shading: 95% confidence interval for the regression line estimated by bootstrap resampling; R and p are Pearson correlation and its p-value, respectively. **g.** Relationship between the strength of excitatory connections from pyramidal onto PV cells and total response similarity. Notation as in panel **f. h–i.** Similarity of trial-average responses but not selectivity for individual visual features predicts the strength of inhibitory connections from PV cells onto pyramidal cells (**h**) and excitatory connections from pyramidal cells onto PV cells (**i**). Error bars are 95% confidence intervals. ***: *p* < 0.001, **: *p* < 0.01, *: *p* < 0.05.

**Supplemental Figure 3:**
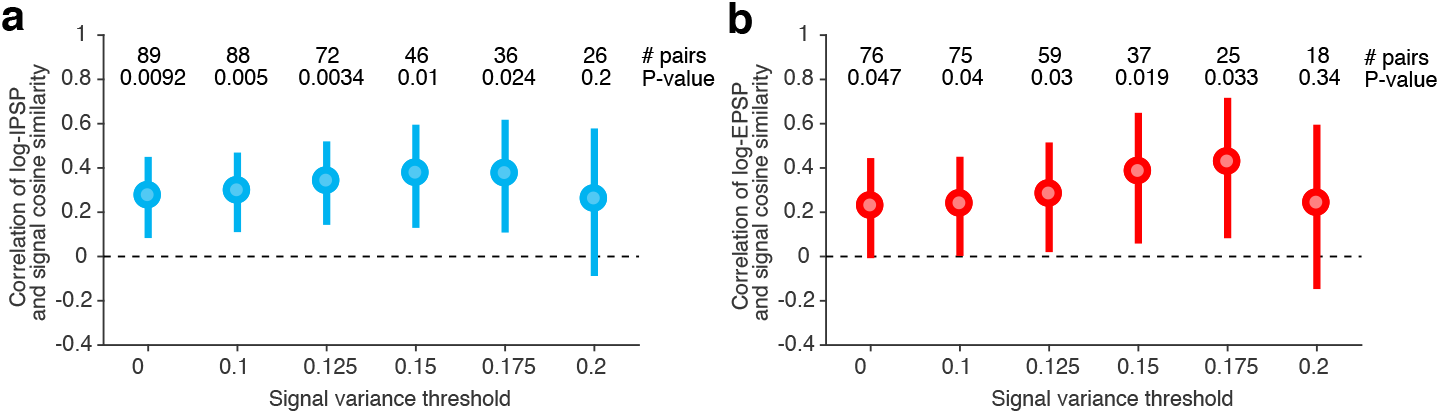
Correlation between signal response similarity and connection strength across a range of responsiveness cut-offs for pyramidal neurons. Error bars are 95% confidence intervals.

Specific inhibition is thought to improve robustness of neuronal tuning to variation in the strength and structure of feedforward inputs^4^, decorrelate spiking of individual neurons^3^, and improve reliability and efficiency of the neural code^2^. On the other hand, broadly tuned inhibition is proposed to sharpen the selectivity of excitatory neurons^21^. To explore the relationship between the rules of excitatory and inhibitory connectivity, and computational properties of cortical networks, we built a spiking neuronal model of recurrent cortical connectivity (Figure 4a). In a simple model configuration, neurons were placed into pre-defined functional cohorts (subnetworks; SN) and preferentially made recurrent synaptic connections within and between subnetworks. Excitatory and inhibitory specificity parameters defined the proportion of total synaptic strength restricted between partners within the same functional cohort; with the remainder of synapses distributed uniformly across the network.

In the presence of uniform, non-specific inhibition, specific excitatory connectivity within subnetworks quickly led to network instability^23^ (Figure 4b-e; stability measure *λ*_*Q*_ > 1 indicates an unbalanced network, see Methods). Unstable network activity modes were caused by recurrent amplification of excitatory activity within subnetworks. Introducing specific connectivity between inhibitory and excitatory neurons caused single inhibitory neurons to become more strongly coupled to subnetworks; this improved network stability by balancing unstable subnetwork activity modes (Figure 4c,f; I Spec. > 0). Conversely, inhibitory specificity that was counter-tuned with excitatory specificity (I Spec. < 0) tended to degrade the stability of a network, by boosting the effectiveness of subnetwork-specific recurrent excitatory connectivity (Figure 4c). The relationship between connection specificity and network stability persisted over a wide range of excitatory and inhibitory connection strengths (Supplemental Figure 7a). Co-tuned inhibitory connectivity therefore permits a greater degree specificity of excitatory connections, while maintaining network stability.

**Figure 4:**
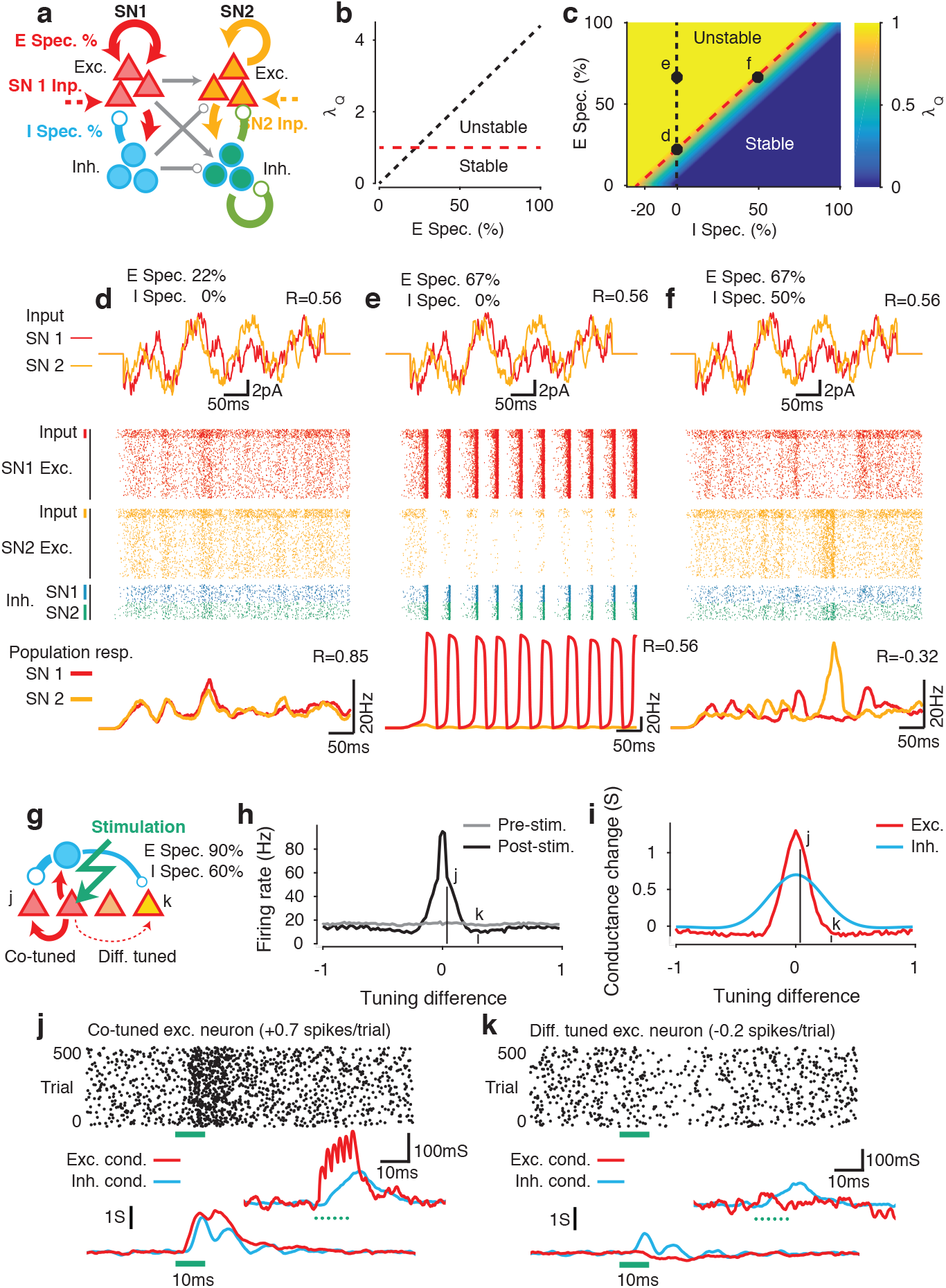
Specific inhibitory connectivity supports network stability and efficient coding. **a.** Two-partition spiking model with excitatory and inhibitory specificity. **b.** In the presence of uniform inhibition (I Spec. =0), specific excitatory connectivity (E Spec) quickly leads to network instability. **c.** Specific inhibitory connectivity (I Spec) balances excitatory specificity, promoting network stability. Dots indicate parameters explored in **d-f**. Black dashed lines in b and c correspond to identical parameters. Red dashed lines indicate the boundary between stable and unstable networks (i.e. *λ* = 1). **d**. A network with E Spec but no I Spec, chosen to be the on the edge of stability. Correlated input currents (top) recruit correlated activity in the network (rasters; middle, and population PSTH; bottom). **e**. An unstable network with strong E Spec and no I Spec. **f.** In a stable network with strong E Spec balanced by moderate I Spec, recruitment and competition lead to strong anticorrelated network activity. **g.** When E and I Spec are smoothly tuned to functional similarity, stimulating a small cohort of excitatory neurons leads to facilitation of co-tuned neurons and suppression in the remainder of the network (**h**). **i-k**. Facilitation (**j**) is driven by strong monosynaptic excitatory inputs, accompanied by strong multi-synaptic inhibitory inputs. Suppression (**k**) is driven by strong multi-synaptic inhibitory inputs in the absence of excitatory inputs. Insets in **j-k** show similar effects evoked by stimulation of a single excitatory neuron.

**Supplemental Figure 7:**
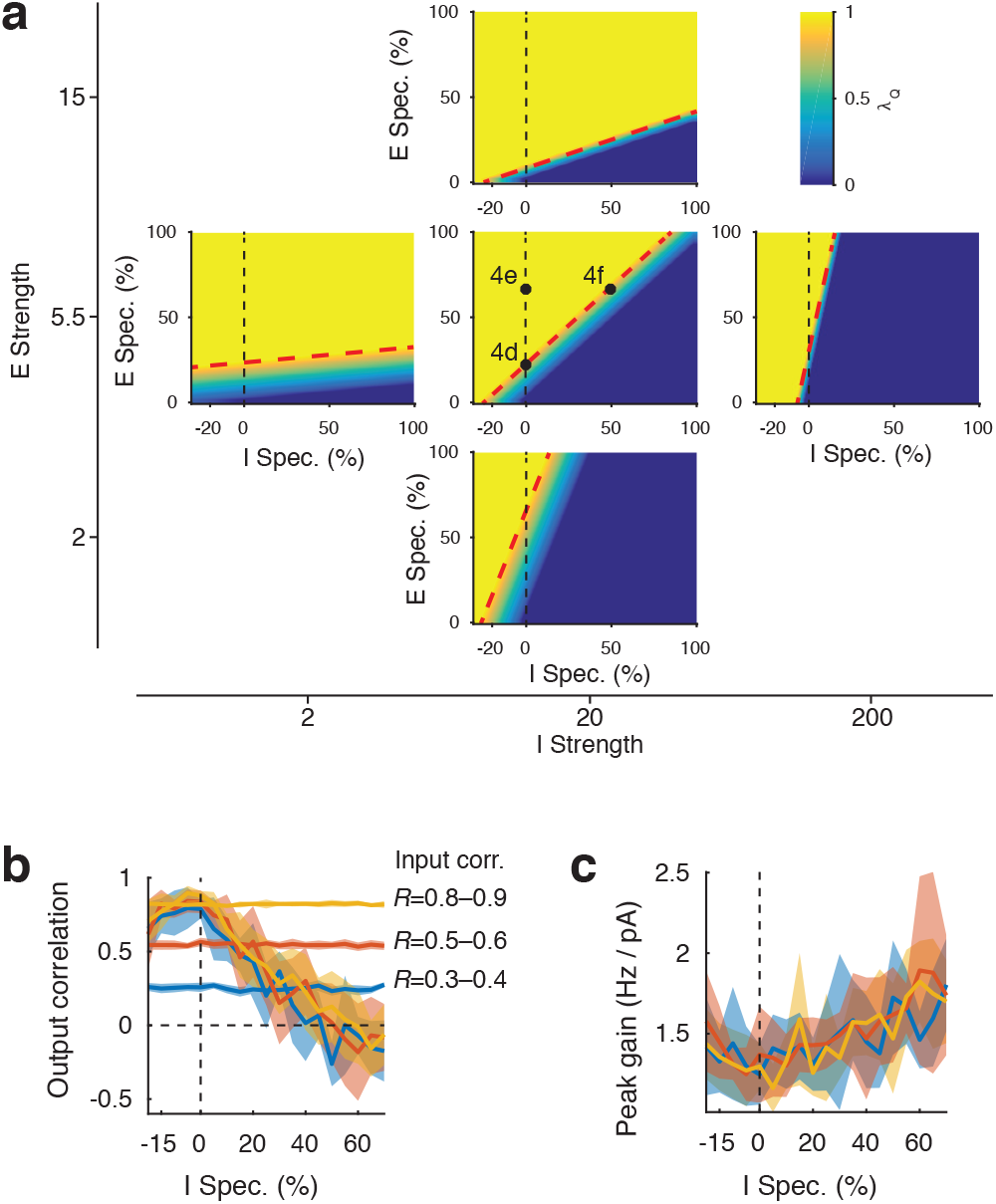
**a.** Specific inhibitory connectivity promotes network stability across a wide range of excitatory and inhibitory connection strength parameters. Dashed black lines indicate I Spec.= 0. Dashed red lines indicate the boundary between stable and unstable networks. **b–c**. Increasing inhibitory and excitatory specificity while maintaining network stability (*λ*_*Q*_ = 0.8) decorrelates subnetwork output independent of input statistics (**b**) and increases output gain (**c**). Shading is standard deviation over different samples of the network input.

How does the interaction between excitatory and inhibitory connection specificity affect the way networks respond to a stimulus? We measured the strength of recruitment and correlation of spiking output of functional cohorts when they were driven with correlated input currents (Figure 4d-f top panels; see Methods). Networks with only weak excitatory connection specificity, not exceeding the threshold of instability, exhibited weak recruitment and highly correlated responses across subnetworks (Figure 4d; correlation *R* = 0.85). Further increasing excitatory specificity while maintaining uniform inhibition lead to network instability and oscillatory bouts of synchronous network activity, as predicted (Figure 4e; correlation *R* = 0.56). On the other hand, networks with specific inhibitory as well as excitatory connections supported strong yet stable recruitment of functional subnetworks but reduced correlations between them (Figure 4f; correlation *R* = −0.32; Supplemental Figure 7b) and increased gain (Supplemental Fig. 7c). This behavior is consistent with the activity of neurons in the cortex, whose mean firing rate correlations are close to zero^3^. Interestingly, the temporal correlations between the responses of the subnetworks were determined by the parameters of specific connectivity in the network and independent of the temporal correlations in the input (Supplemental Figure 7b).

Computational models for competition in cortical networks emphasise global or counter-tuned inhibitory feedback^24–27^. What governs competition and response suppression in the presence of specific inhibitory connectivity as we found in mouse V1? To answer this question we built a model where both inhibitory and excitatory connection strength varied smoothly as a function of tuning similarity (Figure 4g) and stimulated subpopulations of co-tuned excitatory neurons (Figure 4h-k). Neurons with tuning distinct from the stimulated ensemble received strong disynaptic inhibitory input and decreased their firing (Figure 4i,k). Although co-tuned neurons received even stronger inhibitory inputs, they were offset by strong monosynaptic excitation (Figure 4i,j). Thus, specific inhibitory and excitatory connectivity act in concert to facilitate competition between subpopulations of cortical neurons.

We found that connections between inhibitory and excitatory cells in cortex are organized under a similar rule to that of recurrent excitatory connectivity^15^: inhibitory neurons connect more strongly to nearby excitatory neurons with similar responses to visual stimulation. Despite the sea of dense connections between inhibitory and excitatory neurons, selective modulation of connection strength facilitates fine-tuning of inhibition received by individual excitatory neurons and may give rise to the correlations between inhibitory and excitatory synaptic inputs observed in intracellular recordings^5–8^. This organization of inhibitory connectivity provides an circuit substrate for models of cortical networks reliant on feedback inhibition^2,4,28^.

Our results suggest that selective tuning of inhibitory connection strength plays a role in balancing strong recurrent excitatory recruitment, introduced by recurrent connectivity of co-tuned excitatory neurons^15^. Together, the structure of excitatory and inhibitory connections determines the statistical structure of cortical responses, independent of the statistics of feedforward inputs^24^. This circuit architecture ensures that inhibitory and excitatory neurons in mouse visual cortex cooperate to amplify cortical responses while maintaining their sparseness via both disynaptic inhibition across functional subnetworks as well as preferential connectivity within them.

We propose that the history of correlated firing in the network shapes the weights of excitatory and inhibitory connections, which in turn constrain the statistics of cortical responses to feedforward inputs^24^. Since synaptic weights of interneurons and pyramidal cells are determined by their overall response similarity rather than selectivity for individual features of visual stimuli, this wiring rule may be shared by other regions of the cortex.

## Author contributions

S.H., T.M.-F., P.Z., M.H.K., and M.F.I. designed the experiments; P.Z., M.H.K., and M.F.I. carried out the experiments; P.Z., M.H.K., and T.M.-F. analyzed the data; D.R.M. performed the theoretical and modeling work; P.Z., D.R.M., and T.M.-F. wrote the paper.

## Methods summary

All animal procedures were conducted in accordance with institutional animal welfare guidelines and licensed by the Swiss cantonal veterinary office.

Networks of integrate-and-fire spiking neurons with conductance synapse models were simulated using Nest 2.12^29^. Code for reproducing all network simulations is available from figshare^30^. Network stability was quantified by numerically estimating the eigenspectrum of the network weight matrix, under the assumption of linear dynamics.

**Supplemental Figure 4:**
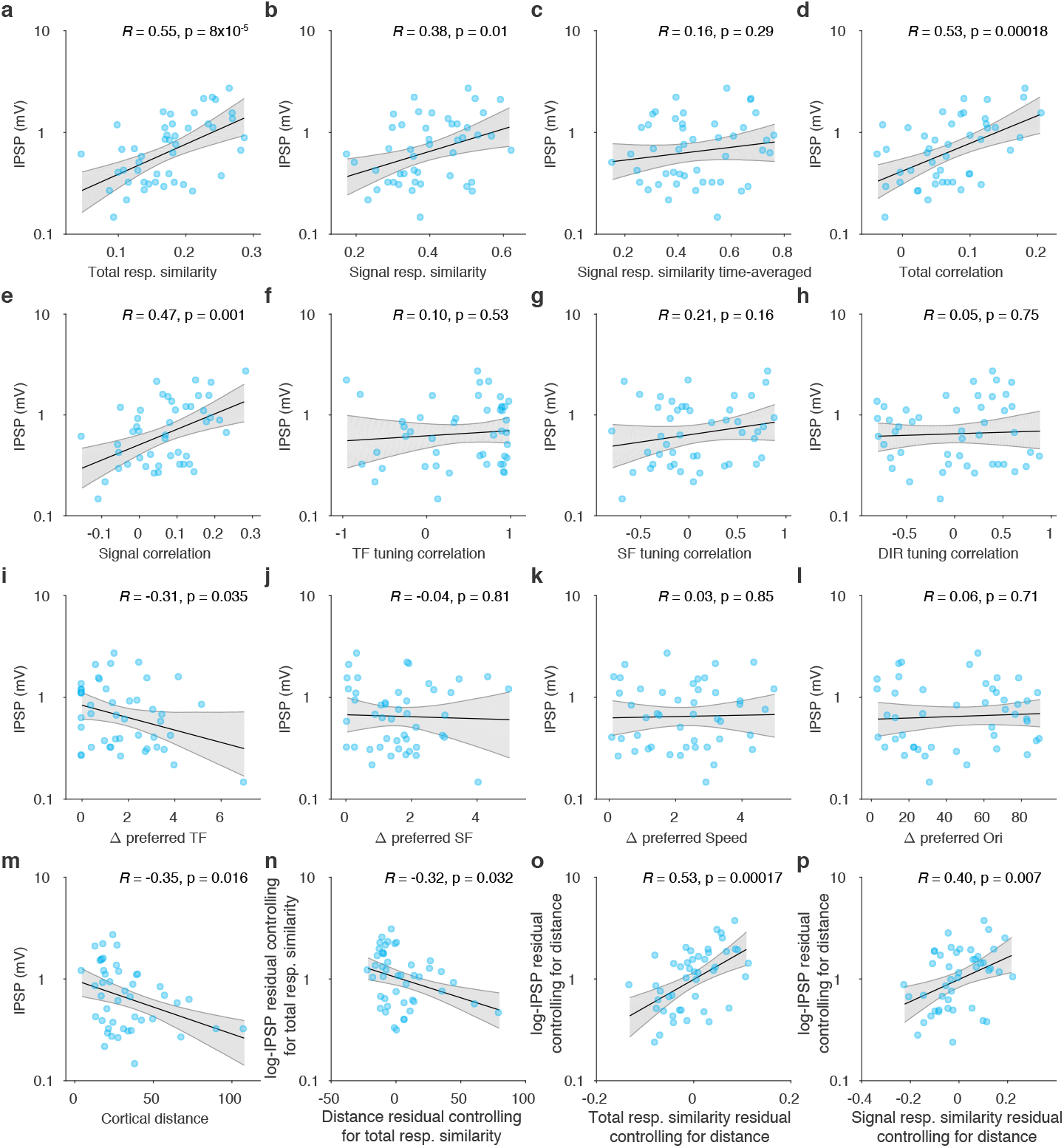
Post-hoc analysis of relationships between IPSP magnitude and the following response similarity metrics. **a**. Total response similarity (as in Figure 3). **b**. Signal response similarity. **c**. Signal response similarity, computed over time-averaged responses to moving phase of the grating. **d**. Total correlation. **e**. Signal correlation. **f**. Correlation of temporal frequency tuning curves, computed from responses to the moving phase of the grating after averaging across spatial frequencies, directions, and time. **g**. Correlation of spatial frequency tuning curves, computed from responses to the moving phase of the grating after averaging across temporal frequencies, directions, and time. **h**. Correlation of direction tuning curves, computed from responses to the moving phase of the grating after averaging across temporal and spatial frequencies, and time. **i**. Absolute difference of preferred temporal frequencies, estimated from model fit. **j**. Absolute difference of preferred spatial frequencies, estimated from model fit. **k**. Absolute difference of preferred speed, estimated from model fit. **l**. Absolute difference of preferred orientations, estimated from model fit. **m**. Distance between cell pairs in the cortical plane. **n**. Residuals of linear regression of log-IPSP and cortical distance against total response similarity. **o**. Residuals of linear regression of log-IPSP and total response similarity against cortical distance. **p**. Residuals of linear regression of log-IPSP and signal response similarity against cortical distance.

**Supplemental Figure 5:**
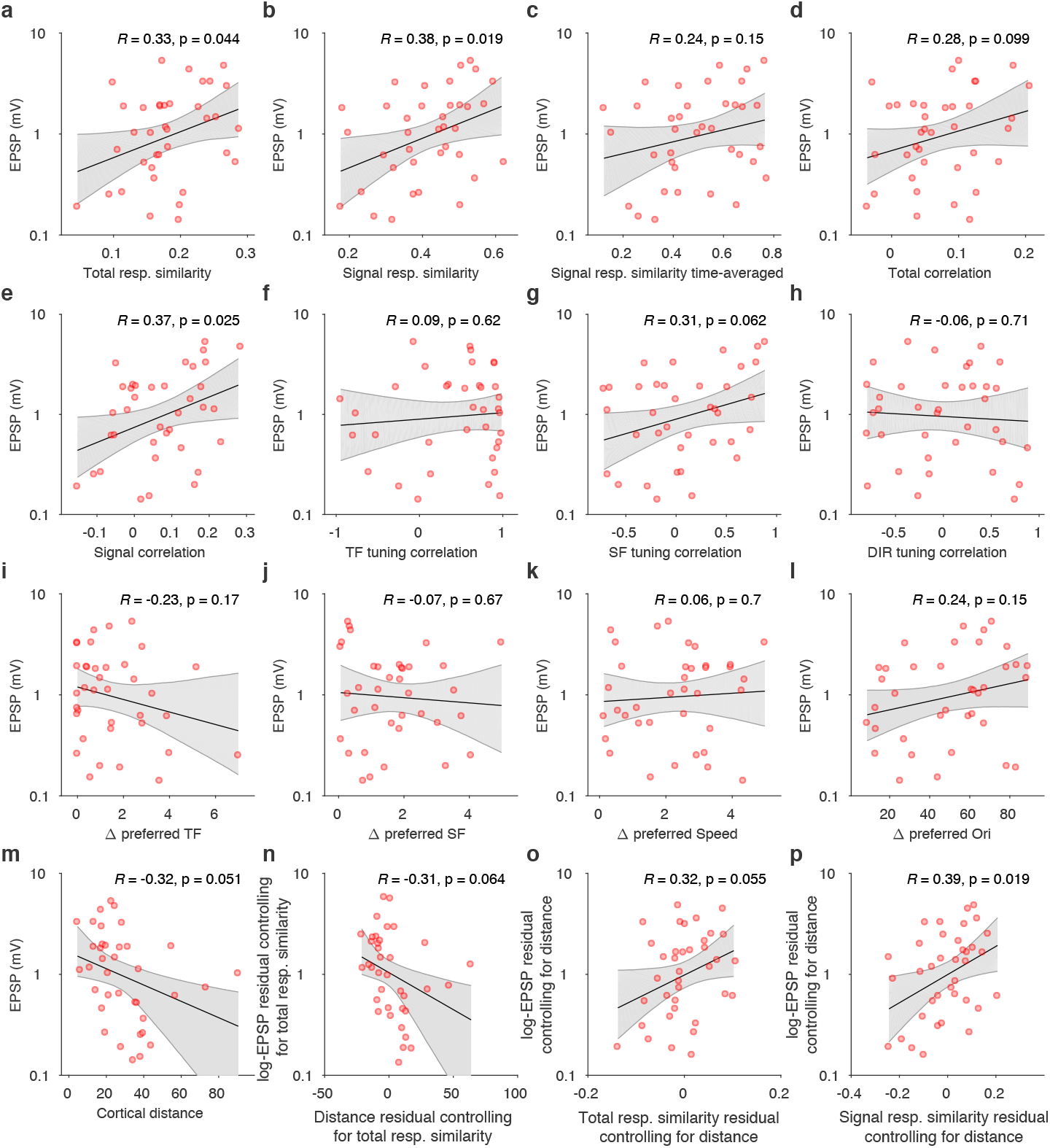
Post-hoc analysis of relationships between EPSP magnitude and individual response similarity metrics.

**Supplemental Figure 6:**
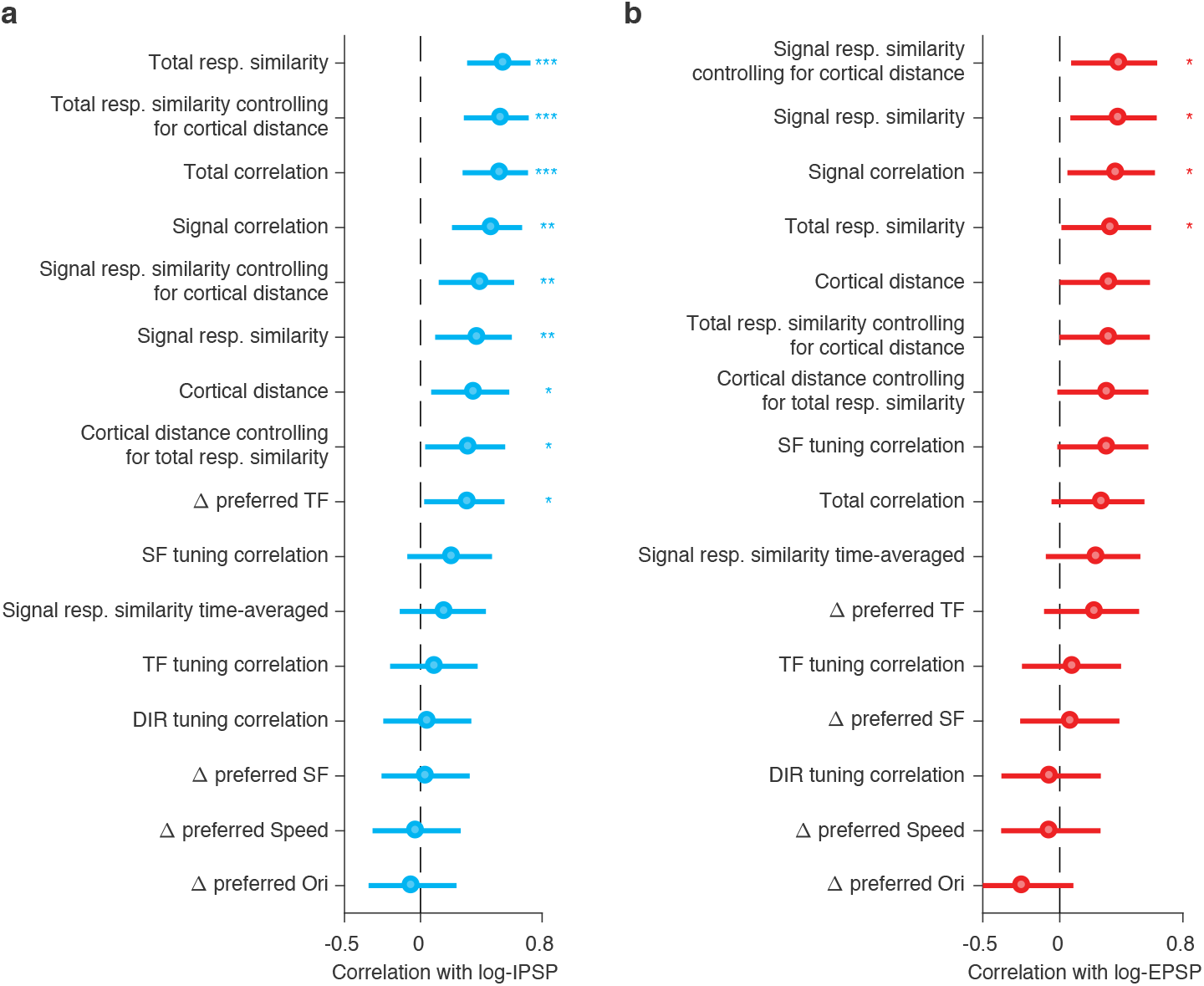
Summary of relationships between connection strength and response similarity metrics from Supplemental Figures 4–5. Error bars are 95% confidence intervals. ***: *p* < 0.001, **: *p* < 0.01, *: *p* < 0.05.

## Full methods

### Responses and connectivity of PV interneurons

#### Animals and surgical procedures

All experiments were conducted in accordance with institutional animal welfare guidelines and licensed by the Swiss cantonal veterinary office. To label parvalbumin expressing interneurons, Ai9 or Ai14 LSL-tdTomato mice were crossed with PV-Cre mice.

At P18–30 male and female animals were anaesthetized with 5 mg/kg midazolam, medetomidine 0.5 mg/kg, and 0.05 mg/kg fentanyl. A metal headplate was implanted exposing the skull over the right visual cortex and 60 nl of AAV 2.1 hSyn-GCaMP6f were injected into V1. To facilitate identification of the injection site *in vitro*, the injection capillary was coated with DiI. Approximately 5 days after the injection, animals were anaesthetized again and a craniotomy 4 mm in diameter was made exposing the visual cortex. A glass coverslip (4 mm diameter, 0.17 mm thickness) was implanted for chronic calcium imaging.

#### In vivo two-photon calcium imaging

Awake mice were head-fixed and allowed to run on a styrofoam wheel. A monitor (47 cm wide) was placed 20–25 cm away from the eye spanning a field of view of 122 visual degrees. Monitor position was adjusted such that the center of the screen matched the preferred retinotopic location of the imaging site, as judged by two-photon fluorescence responses to grating patches flashed at different locations on the screen.

Fluorescence signals were recorded using a ThorLabs Bergamo and a ThorLabs Bergamo II resonant scanning two-photon microscopes with a Nikon 16x water-immersion objective (NA 0.8) operated using ScanImage 4 or 5.1 software. GCaMP6f fluorescence was imaged using 930 nm excitation at 10–30 mW with a 520/40 nm emission filter (Chroma). Volumes of 8 frames spanning 80 *µ*m in depth and ~350–520 *µ*m in X-Y were acquired at 3.75 Hz using a piezo focuser (PI P-726). For identification of PV-positive neurons, tdTomato fluorescence was imaged at 930 nm with a 607/70 nm emission filter (Semrock).

To prevent the light from the monitor from interfering with imaging, the monitor backlight was controlled by a custom electronic circuit and only switched on during the turn-around of the resonant X mirror^31^. Gratings of 6 spatial frequencies, 6 temporal frequencies and 8 directions were interleaved randomly and presented without gaps. Each grating first remained stationary for 1.3 or 2.1 seconds (5 or 8 volumes), before moving for 2.1 seconds (8 volumes). A single presentation of the stimulus set constituted a single imaging segment, 6–10 of which were repeated during each imaging session.

#### In vivo/*in vitro* registration

Following the in vivo imaging session, animals were anaesthetized using isolfuorane (1.5–2%). A detailed Z-stack of GCaMP6f, tdTomato, and DiI fluorescence at the imaging site was acquired at 830 nm, spanning the volume from 0 to ~300 *µ*m below pia. Since GCaMP6f fluorescence at this wavelength does not depend on calcium concentration, neurons could be readily identified independent of their level of activity.

The slice containing the *in vivo* imaging site was identified *in vitro*, using DiI fluorescence and parenchymal blood vessels as landmarks. As *in vivo*, a Z-stack was acquired at 830 nm spanning all or most of the slice thickness. *In vivo* and *in vitro* volumes were then aligned using custom software written in Matlab (https://github.com/znamensk/RegisterStack). Four or more manually selected control points were used to estimate forward (*in vivo to in vitro*) and inverse (*in vitro to in vivo*) affine transformation matrices using least squares regression. The quality of the registration was verified by rotating and reslicing the *in vivo* Z-stack in the coordinate frame of the *in vitro* volume using the forward transformation. If necessary, control points were added or replaced until the registration quality was satisfactory. A similar procedure was used to register the *in vivo* Z-stack with the *in vivo* functional imaging planes.

The inverse transformation matrices were used to identify the *in vivo* imaging ROI corresponding to each of the neurons recorded *in vitro*. Registration accuracy was confirmed by manual inspection and 12/250 patched neurons, which were located within the imaging volume but could not be unambiguously identified, were excluded from further analysis.

#### In vitro whole-cell patch clamp recording

At least 12 hours after the imaging session, mice were lightly anesthetized with sodium pentobarbital and transcardially perfused with a cold choline chloride based solution containing (in mM): 110 choline chloride, 25 NaHCO_3_, 25 D-glucose, 11.6 sodium ascorbate, 7 MgCl_2_, 3.1 sodium pyruvate, 2.5 KCl, 1.25 NaH_2_PO_4_, and 0.5 CaCl_2_^32^ with ~325 mOsm. Visual cortex slices (300–350 *µ*m thickness) were cut coronally on a vibrating blade microtome (VT1200S, Leica Biosystems) with the same choline chloride based solution bubbled with 95% O_2_/5% CO_2_. Then, the brain slices were incubated at 34°C for 20–40 min with artificial cerebrospinal fluid (ACSF) solution containing 125 mM NaCl, 2.5 mM KCl, 1 mM MgCl_2_, 1.25 mM NaH_2_PO_4_, 2 mM CaCl2, 26 mM NaHCO3, 25 mM D-glucose; 315–320 mOsm adjusted by adding the amount of D-glucose, bubbled with 95% O_2_/5% CO_2_, pH 7.4. Afterwards, the brain slices were continuously maintained at room temperature before being transferred to the recording chamber.

In vitro imaging and patch-clamp recordings were performed with Scientifica MP-1000 multiphoton imaging microscope and a mode-locked Ti:sapphire laser (Vision-S, Coherent) with a Nikon 16x water-immersion objective (NA 0.8). Scanning and image acquisition were controlled by SciScan (Scientifica) and custom software written in LabVIEW (National Instruments).

Recording pipettes were mounted on remote-controlled motorized micromanipulators (MicroStar, Scientifica). Recording pipettes were made using thick-walled filamentous borosilicate glass capillaries (G150F-4, Warner Instruments) using a horizontal puller (P-1000, Sutter Instrument) adjusted to produce pipette resistance of 7–8 MOhm with a long taper when filled with intracellular solution in 34°C ACSF. The potassium based internal solution containing 5 mM KCl, 115 mM K-gluconate, 10 mM HEPES, 4 mM Mg-ATP, 0.3 mM Na-GTP, 10 mM Na-phosphocreatine, 0.1% w/v Biocytin; osmolarity 290–295 mOsm, pH 7.2 was used. Liquid junction potentials were not corrected.

Whole-cell patch-clamp recordings were carried out at ~34°C using Multiclamp 700B amplifiers (Axon Instruments) and custom-written MATLAB software (MathWorks). Up to 6 cells, at least one of which was a tdTomato-expressing PV neuron, were targeted simultaneously. All tdTomato positive cells showed fast-spiking firing profiles during current injection and evoked inhibitory post-synaptic potentials in connected pyramidal cells. Pyramidal neurons were identified based on their regular spiking firing profiles. During 9 out of 35 experiments, the experimenter could target cells based on the correlation of their calcium traces in the in *vivo* imaging dataset.

To reveal inhibitory inputs, pyramidal cells were depolarized to −55–50 mV by current injection. To test for the presence of synaptic connections, two or five presynaptic spikes were evoked by current injections at 30 Hz in each cell sequentially repeated 20–150 times, while searching for corresponding postsynaptic responses. Recordings with postsynaptic cell series resistances below 35 MOhm were included for analysis.

#### Data analysis: preprocessing

Two-photon imaging frames were registered using a phase-correlation algorithm. ROIs were identified based on the mean fluorescence image and their fluorescence time series were extracted. To correct for bleaching and other sources of non-stationarity, fluorescence traces were high-pass filtered at 0.0019 Hz. The neuropil fluorescence was measured in ~40 *µ*m radius around the center of each cell, excluding any detected ROIs.

Since the amount of neuropil signal within the ROI depends on window quality, imaging depth and axial position of the imaging plane within the cell, we used robust linear regression (MATLABs *robustfit* with the default bisquare weight function) to estimate the coefficient of neuropil contamination *b*_*neuropil*_ for each cell. This procedure allowed us to estimate the amount of neuropil contamination for sparsely active ROIs. However, for densely active cells, which were often highly correlated with the neuropil signal, robust regression could overestimate the degree of neuropil contamination and return correction coefficients exceeding 1. In this case, to avoid overcorrecting we fixed *b*_*neuropil*_ to 0.7. This procedure was carried out separately for each imaging session segment, containing a single presentation of the stimulus set. The cell’s corrected fluorescence signal was calculated as:

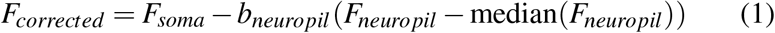

To calculate *dF/F*_0_, we fit a mixture of two Gaussians to the distribution of neuropil-corrected fluorescence values and used the mean of the smaller Gaussian as an estimate of baseline fluorescence *F*_0_.

#### Data analysis: responsiveness and selectivity

To identify robustly responsive neurons without making assumptions about the shape of the neurons tuning curve or the time-courses of their responses, we computed the fraction of variance explained by the visual stimulus:

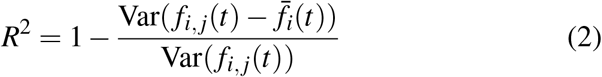

where *f*_*i,j*_(*t*) is the dF/F of the neuron during frame *t* of the *i*^th^ stimulus type on the *j*^th^ trial, and

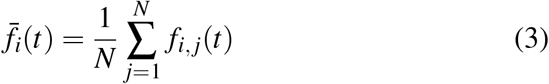

is the trial average response on frame *t* of the *i*^th^ stimulus type, and *N* is the number of trials.

We classified neurons as responsive, if the visual stimulus explained > 15% of the variance of their dF/F responses. Note that while this measure is related to the F-statistic used in ANOVA, our criterion is more stringent than an F-test. While 198/250 ROIs studied in the in vivo/in vitro experiments were significantly responsive by one-way ANOVA (*p <* 0.05), only 151/250 of them passed our variance explained threshold.

To compute the cells selectivity we first computed the mean dF/F response during the moving phase of the grating (*a*^th^ to *b*^th^ frame) for the each stimulus type:

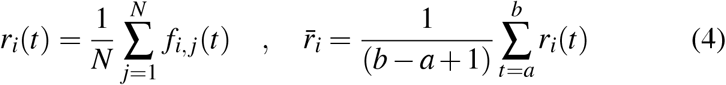

Cells’ selectivity was defined as the skewness of 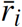:

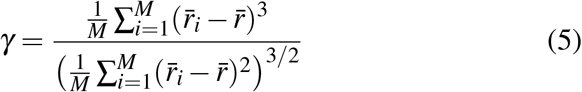

where 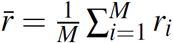 is the mean response across all stimulus types, and *M* is the number of stimulus types.

**Table 1:** Parameters of the direction, spatial and temporal frequency fit.

| Parameter | Description |
| --- | --- |
| $b$ | Response offset |
| $r_{max}$ | Response at preferred spatial, temporal frequency, and direction, $0 < r_{max} \leq 2 \cdot \max(r_{i,j})$ |
| $SF_{pref}, TF_{pref}$ | Preferred spatial and temporal frequency in octaves, constrained to lie within 1 octave of the range probed in the experiment |
| $\alpha$ | Orientation of the SF/TF tuning Gaussian, $0^\circ \leq \alpha \leq 90^\circ$ |
| $\sigma_x, \sigma_y$ | Tuning width of SF/TF Gaussian, constrained between 0.25 and 4 octaves |
| $\sigma_{Dir}$ | Width of direction tuning, $0^\circ \leq \sigma_{Dir} \leq 180^\circ$ |
| $q$ | Relative magnitude of the response to the null direction, $0 \leq q \leq 1$ |
| $\theta_{pref}$ | Preferred direction, $0^\circ \leq \theta_{pref} \leq 360^\circ$ |

#### Data analysis: direction and SF/TF tuning

To characterize the direction, spatial and temporal frequency tuning of individual cells, we fit their single trial responses during the moving phase of the grating

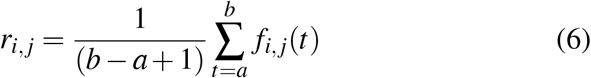

as the product of a double Gaussian in direction space and a 2D Gaussian in spatial and temporal frequency space with arbitrary orientation. Parameters of the fit are described in Table 1. Let *x*_*i*_ and *y*_*i*_ be defined as:

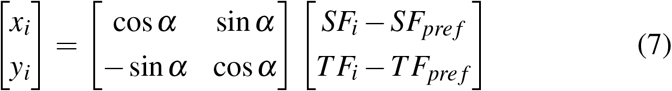

The response *r*_*i,j*_ was then modeled as:

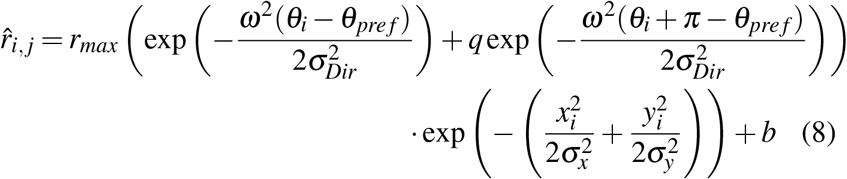

where *ω*(*θ*) wraps angles onto the interval between 0 and *π*:

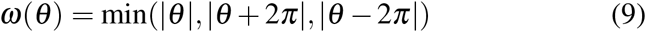

We determined the values of the fit parameters using *lsqnonlin* in MAT-LAB by minimizing 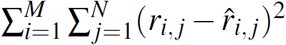 with the constraints specified in Table 1.

*FWHM*_*Dir*_ was calculated as 2.355 ⋅ *σ*_*Dir*_. If *FWHM*_*Dir*_ exceeded 180 degrees, cells were classified as untuned for direction/orientation and excluded from direction selectivity analysis Figure 1d. *FWHM*_*SF*_ and *FWHM*_*TF*_ were calculated from tuning curves derived by estimating 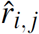 using Eq. 8 with the estimated fit parameters over a range of spatial and temporal frequencies (0.0025 to 2.56 cycles per degree, and 0.125 to 16 Hz). If *FWHM*_*SF*_ or *FWHM*_*TF*_ exceeded 6 octaves, cells were classified as untuned for spatial or temporal frequency, respectively.

#### Data analysis: population pooling

Pooled population responses were calculated by summing the activity of neurons within the imaging volume weighted by their proximity to the cell of interest. To facilitate comparisons between predictions based on the cells’ own responses and population activity, we first computed the mean response for each cell, reserving one trial at a time:

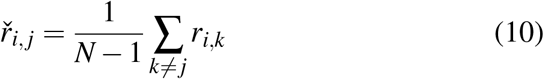

The limit on the prediction of single cells’ tuning, imposed by trial-to-trial variability of their responses, was estimated as the mean Pearson correlation coefficient of single trial responses *r*_*i,j*_ and 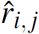:

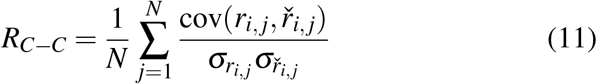

To compute the population prediction 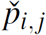, we summed the normalized responses of each cell, scaled by a Gaussian function of their distance to the cell of interest with a specified length scale and excluding cells separated by < 10 *µ*m in the same or adjacent imaging planes to avoid fluorescence cross-talk:

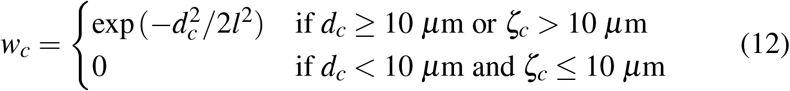

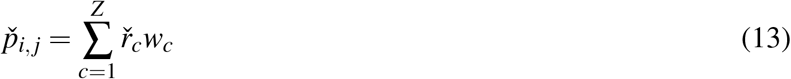

where 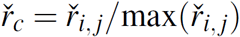 is the normalized response of cell *c* excluding trial *j*, *d*_*c*_ is cortical distance between cell *c* and the cell of interest and *ζ*_*c*_ is the separation of their imaging planes, *l* is the spatial scale of pooling, and *Z* is the number of cells in the imaging volume. Performance of the population prediction was quantified as the Pearson correlation coefficient of *r*_*i,j*_ with 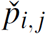 and averaged across trials:

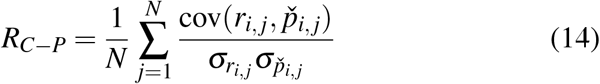

To compute population selectivity, we first calculated the mean population response across all trials:

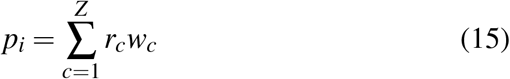

where 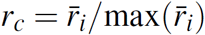 is the normalized trial-average response of cell *c* (See Eq. 4). Population selectivity was then calculated as the skewness of *p*_*i*_:

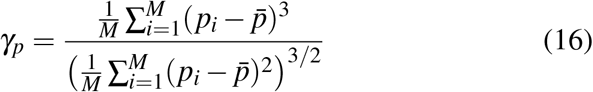

where 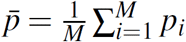 is the mean population response across all stimulus types.

#### Data analysis: synaptic connectivity

Connected pairs were identified by inspecting all of the acquired postsynaptic traces. However, as the magnitude of postsynaptic responses often fluctuated following repeated presynaptic stimulation, we limited our analysis of the magnitude of postsynaptic responses to at most the first 20 trials. After subtracting the baseline membrane potential (average Vm in 5 ms preceding presynaptic stimulation), postsynaptic responses were aligned to the time of the presynaptic spike and averaged. Cell pairs where the presence or absence of a connection could not be unambiguously determined were excluded from further analysis. IPSP and EPSP magnitudes were estimated as the minimum or maximum of the mean postsynaptic trace, respectively. In 10/124 PV to pyramidal cell pairs and 5/120 pyramidal to PV cell pairs, where presynaptic stimulation evoked multiple spikes and PSPs overlapped, IPSP and EPSP magnitudes were estimated in the window preceding the onset of the second PSP. Excluding these pairs from the analysis did not affect our conclusions.

As IPSP and EPSP magnitudes roughly followed a log-normal distribution, we used log-transformed PSP magnitudes to quantify their relationship with each other, as well as response similarity and other stimulus features.

We used the following procedure to test whether the correlation between IPSP and EPSP magnitudes for reciprocally connected Pyr-PV pairs could be explained by variation in slice quality. We first selected PV cells, for which we measured the strength of reciprocal connections with at least 2 pyramidal cells. We then normalized the log-IPSP and log-EPSP magnitudes by their mean for each such PV cell:

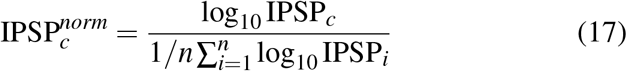

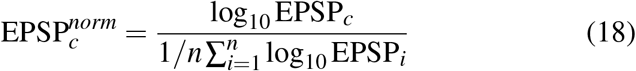

where IPSP_*c*_ and EPSP_*c*_ are the magnitudes of connections between the PV cell and the *c*^th^ pyramidal cell. We quantified the relationship between normalized IPSP and EPSP magnitudes across all recordings as the Pearson product moment correlation coefficient:

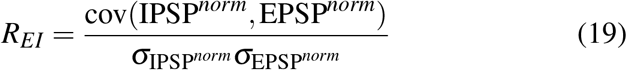

To assess the statistical significance of this relationship, we randomly permuted IPSP magnitudes for each PV cell, destroying any relationship between IPSP and EPSP strength, and recalculated the correlation coefficient. The P-value was estimated as the proportion of reshuffled samples, where the absolute value of the correlation exceeded the one observed empirically.

#### Data analysis: relating synaptic connectivity and in vivo responses

Cell pairs profiled in vitro were only included in the analysis in Fig. 3 if both cells were located within the imaging volume and could be reliably identified. Of the 157 PV cell to pyramidal cell connections and 164 pyramidal cell to PV cell connections, 118 and 116 respectively passed this criterion. Since fluorescence cross-talk between overlapping ROIs would inflate our estimate of response similarity, we excluded cell pairs separated by < 10 *μ*m in the same or adjacent imaging planes. Finally, we restricted our analysis to pairs where the pyramidal cell was visually responsive and passed the signal variance threshold of 15%. This dataset included 56 PV cell to pyramidal cell pairs (from 21 mice) and 55 pyramidal cell to PV cell pairs (from 20 mice), of which 46 (from 18 mice) and 37 (from 19 mice) respectively were connected.

To compute total response similarity between pairs of ROIs, we computed the dot product of their *dF*/*F*_0_ traces normalized by the product of their norms for each imaging segment, and used the mean of these values:

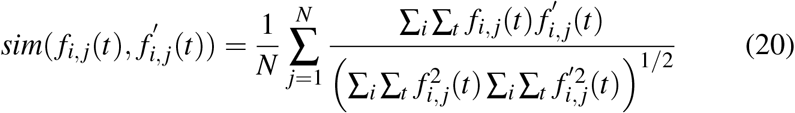

where *f*_*i,j*_(*t*) and 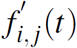 are *dF/F*_0_ traces of the two cells for the *i*^th^ stimulus type on the *j*^th^ trial. To compute signal response similarity, we used the trial-average responses *r*_*i*_(*t*) and 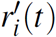, as defined in Eq. 4:

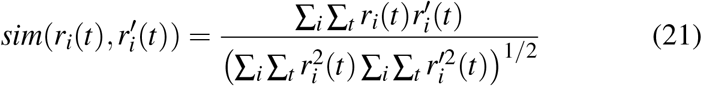

Tuning similarity metrics for individual visual features were computed as follows:

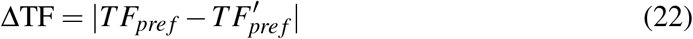

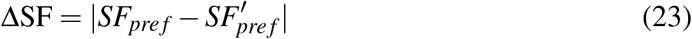

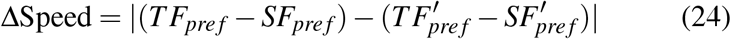

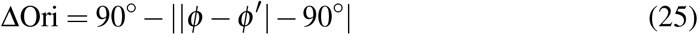

where *ϕ* and *ϕ*’ are the preferred orientations of each cell, *θ*_*pref*_ mod 180°.

Additional response similarity metrics in Supplemental Figures 4-6 were computed as follows:

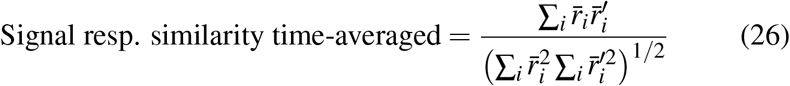

**Table 2:** **Parameters used in the network models.** Other parameters as described in the methods text.

| Parameter | Description |
| --- | --- |
| $N$ | Number of neurons in the network ($N = 5000$ for partitioned model; $N = 1000$ for feature tuning model) |
| $N_S$ | Number of subnetwork partitions in partitioned model ($N_S = 2$) |
| $w_E, w_I$ | Total output synaptic weight made by single excitatory and inhibitory neurons, respectively ($w_E, w_I > 0$) |
| $f_E, f_I$ | Proportion of excitatory and inhibitory neurons in the network ($f_E + f_I = 1$; $f_I = 20\%$) |
| $d_{IE}$ | Factor defining how much stronger $E \rightarrow I$ synaptic connections are compared with $E \rightarrow E$ ($d_{IE} \geq 1$; $d_{IE} = 2$) |
| $s_E$ | Proportion of recurrent excitatory synapses that are restricted to be made within the same subnetwork partition ($0 \leq s_E \leq 1$) |
| $s_I$ | Proportion of synapses between excitatory and inhibitory neurons, and recurrent inhibitory synapses, that are restricted to be made within the same subnetwork partition ($0 \leq s_I \leq 1$) |
| $\kappa$ | Concentration parameter controlling synaptic specificity over feature parameter in feature tuning model, for neurons of class $A$ targetting neurons of class $B$ ($\kappa_{EE} = 16$; $\kappa_{EI}, \kappa_{IE}, \kappa_{II} = 4$) |

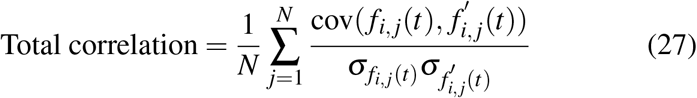

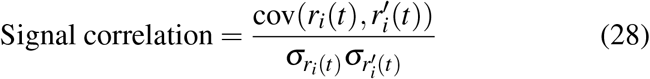

SF, TF, and direction tuning correlations were each computed in a similar manner, after first averaging 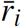 across the other two stimulus dimensions.

The relationship between response similarity and connection probability was quantified using logistic regression. The relationship between response similarity or other response metrics and PSP magnitude was quantified as the Pearson product moment correlation coefficient of the functional similarity metric and log-transformed PSP magnitude.

Partial correlations controlling for distance were calculated using *partialcorr* in MATLAB, computing the Pearson correlation coefficient of residuals of linear regression of similarity metrics and log-PSP magnitude against distance.

### Models of specific inhibitory and excitatory connectivity

Code to generate, simulate and analyse all models described in this paper is available online^30^.

#### Partitioned network model

Partitioned networks were generated and analysed as described previously^23^. Parameters used to build the network models are defined in Table 2. Briefly, a network of *N* neurons was defined to contain *N* ⋅ *f*_*E*_ and *N* ⋅ *f*_*I*_ excitatory and inhibitory neurons. In this paper, dense connections were made between all potential partners (i.e. synaptic fill factors *h*_*E*_,*h*_*I*_ = 1). Networks were partitioned into equal sized subnetworks, with both excitatory and inhibitory neurons assigned a subnetwork identity. The strength of synaptic connections between neurons was modulated by whether pairs of neurons were members of the same or different subnetworks.

To generate connections between neurons we defined specificity parameters *s*_*E*_ and *s*_*I*_, which determined what proportion of output synapses from single neurons were made according to specific connection rules. A proportion *s*_*E*_ of total recurrent excitatory synaptic weight was reserved to be made with partners in the same subnetwork. The remainder of recurrent excitatory weight was distributed uniformly over the network. Similarly, a proportion *s*_*I*_ of total recurrent inhibitory weight was reserved to be made within the same subnetwork. The same parameter *s*_*I*_ modulated specific connectivity between excitatory and inhibitory neurons. In these models we assumed that *E → E* and *E → I* synaptic specificity was equal.

Output weights for all neurons were normalized such that the total output weights were equal to *w*_*E*_ and *w*_*I*_ for excitatory and inhibitory neurons respectively, distributed proportionally across excitatory and inhibitory targets according to *f*_*E*_ and *f*_*I*_.

The synaptic weights *w*_*BA*_ between two neurons in the same subnetwork from class *A* to class *B* were therefore given by

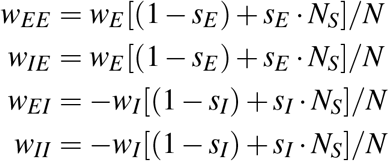

and between neurons in different subnetworks given by

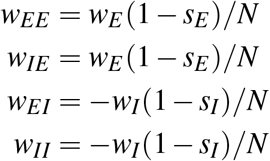

#### Visual feature tuning model

To examine the effect of smooth functional relationships between neurons on connectivity and network computation, we defined a richer model where connections between excitatory and inhibitory neurons were governed by similarity of tuning to visual features. Since no single visual feature was found to explain the entirety of synaptic connection specificity in mouse V1, we defined connectivity over an arbitrary visual feature tuning parameter. Excitatory and inhibitory neurons comprised 80% and 20% of the neuronal population respectively, as for the partitioned networks above. Neurons were assigned a preferred tuning value *γ*, uniformly distributed over (0,1]. Synaptic connection strength between two neurons *i* and *j* was modulated by similarity measured over this tuning parameter. The connection probability from a neuron *i* of class *A* to a neuron *j* of class *B* was given by the circular function

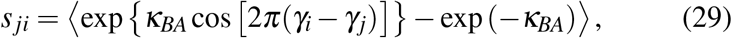

where *κ*_*BA*_ is a concentration parameter modulating connections from class *A* to class *B.* The notation 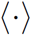 indicates the expression within the brackets has been normalised to define a probability density function over target synaptic partners. The values *s*_*ji*_ were composed into the *N* × *N* matrix

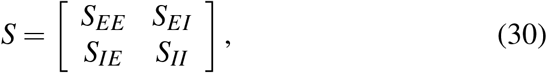

with the blocks *S*_*BA*_ defining the connections from class *A* to class *B.* The synaptic weights connecting neurons were also modulated by specificity parameters, defining a proportion of synapses that were made under the function in Eq.29. The remainder of synapses were distributed uniformly over the network. The weight matrix for the network was then given in block form by

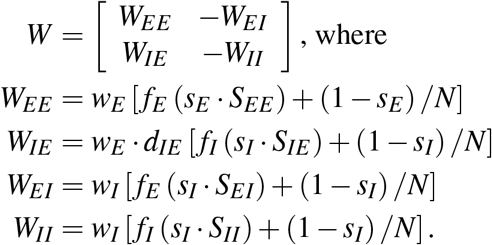

#### Network input

Each neuron in the network received a barrage of independent Poisson input spikes at 2400 Hz, designed to mimic ongoing spontaneous activity in cortex. External input was provided by injecting currents into the excitatory population alone, since input provided directly to the inhibitory population can give rise to suppression of excitatory activity through feed-forward inhibition. Since our goal was to examine recurrent computations, we wished to exclude this feed-forward effect from our analysis.

For the partitioned network models, white noise input currents were generated with standard deviation *σ* = 1 over 1 ms, then smoothed with a box-car filter of width 50 ms. The currents were subsequently normalized to have a peak to peak amplitude of 10 pA, and shifted to have a mean of 15 pA. A single white noise current was generated for each partition; this procedure was repeated in a brute-force approach until a set of input currents was found with a desired correlation coefficient *R.* Each resulting white noise current was subsequently injected into a subset of 12.5% of the excitatory population of each subnetwork.

For the feature tuning model, perturbation of small cohorts of neurons (Fig. 4g–k) was performed by first increasing background activity to 3500 Hz per neuron. 21 excitatory neurons with similar feature tuning *γ* were then stimulated by injecting currents of 100 pA (corresponding to 2.7% of the excitatory network). Perturbation of a single excitatory neuron was performed by injecting a current of 0.2 *μ*A (insets in Fig. 4j,k).

#### Linear stability analysis

To estimate the stability of the networks described above, we examined a non-spiking network model with dynamics governed by

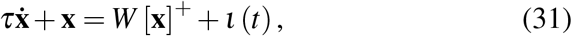

where 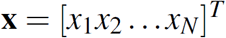 is a vector of neuron activations, *I*(*t*) is an external input provided to the network, τ is a lumped neuron time constant and [⋅]^+^ is the linear-threshold activation function [*x*]^+^ = max(*x*,0). Under this formulation, the eigenspectrum of the weight matrix *W* provides information about the stability of various network activity patterns, as a function of the network connectivity parameters^23^. We examined the stability of the networks in the state where all neurons were active (i.e. 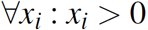).

For the two-partition network, the eigenvalues related to subnetwork partitioning in the presence of excitatory and inhibitory specificity are given by the closed-form equation

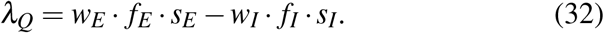

For the feature tuning model, the eigenspectra of the weight matrices were estimated numerically.

#### Spiking simulations

Simulations of integrate and fire spiking neurons with alpha-function conductance-based synapses were performed using Nest 2.12^29^ and Matlab (Mathworks). Networks were connected following the weight matrices defined above. Weights were scaled such that recurrent spiking dynamics matched those of a linear network as closely as possible. That is, a recurrent excitatory weight of ‘1’ placed a self-connected neuron in a critical regime on the edge of stability.

